# A TetR-family protein activates transcription from a new promoter motif associated with essential genes for autotrophic growth in acetogens

**DOI:** 10.1101/668756

**Authors:** Renato de Souza Pinto Lemgruber, Kaspar Valgepea, Ricardo Axayacatl Gonzalez Garcia, Christopher de Bakker, Robin William Palfreyman, Ryan Tappel, Michael Köpke, Séan Dennis Simpson, Lars Keld Nielsen, Esteban Marcellin

## Abstract

Acetogens can fix carbon (CO or CO_2_) into acetyl-CoA via the Wood-Ljungdahl pathway (WLP) that also makes them attractive cell factories for the production of fuels and chemicals from waste feedstocks. Although most biochemical details of the WLP are well understood and systems-level characterisation of acetogen metabolism has recently improved, key transcriptional features such as promoter motifs and transcriptional regulators are still unknown in acetogens. Here, we use differential RNA-sequencing to identify a previously undescribed promoter motif associated with essential genes for autotrophic growth of the model-acetogen *Clostridium autoethanogenum*. RNA polymerase was shown to bind to the new promoter motif using a DNA-binding protein assay and proteomics enabled the discovery of four candidates to potentially function directly in control of transcription of the WLP and other key genes of C_1_ fixation metabolism. Next, *in vivo* experiments showed that a TetR-family transcriptional regulator (CAETHG_0459) and the housekeeping sigma factor (σ^A^) activate expression of a reporter protein (GFP) in-frame with the new promoter motif from a fusion vector in *E. coli*. Lastly, a protein-protein interaction assay with the RNA polymerase (RNAP) shows that CAETHG_0459 directly binds to the RNAP. Together, the data presented here advance the fundamental understanding of transcriptional regulation of C_1_ fixation in acetogens and provide a strategy for improving the performance of gas-fermenting bacteria by genetic engineering.

## 1. Introduction

The Wood-Ljungdahl pathway (WLP) of acetogens is speculated to be the first biochemical pathway on Earth that emerged when the atmosphere was still highly reduced and rich in CO, CO_2_, and H_2_ (Fuchs, 2011; Russell and Martin, 2004; Weiss et al., 2016). These C1 gases can be converted into acetyl-CoA through the WLP (Ragsdale and Pierce, 2008; Wood, 1991) and acetogens are the only known organisms using the WLP as a terminal electron-accepting, energy-conserving process to fix CO_2_ into biomass (Drake et al., 2006; Fuchs, 2011). This pathway is responsible for the production of acetic acid in quantities surpassing the billion ton mark annually. It is estimated that the pathway contributes to fixing ~20% of the CO_2_ on Earth (Drake et al., 2006; Ljungdahl, 2009). All this takes place with the WLP operating at the edge of thermodynamic feasibility (Schuchmann and Müller, 2014) and requires the use of the third mode of energy conservation, electron bifurcation, which likely contributed to the emergence of life on Earth (Herrmann et al., 2008; Li et al., 2008; Nitschke and Russell, 2011). Acetogens are also attractive cell factories for the sustainable production of fuels and chemicals from gaseous waste feedstocks (e.g. syngas from gasified municipal solid waste and industrial waste gases) (Claassens et al., 2016; Dürre and Eikmanns, 2015; Liew et al., 2016; Molitor et al., 2016). While the field has advanced enourmously in the last decade (Liew et al., 2016; Molitor et al., 2016), better fundamental understanding of acetogen metabolism is needed to guide rationale metabolic engineering, for example, to increase their substrate uptake or product yields.

Recent quantitative studies of acetogen physiology have expanded understanding of their metabolism considerably (reviewed in (Molitor et al., 2017; Schuchmann and Müller, 2014)). Although most biochemical details of the WLP are well established (Ragsdale, 1991, 1997, 2008) and systems-level understanding of acetogen metabolism has recently improved (Valgepea et al., 2017a, 2018), key transcriptional features such as promoter motifs and transcriptional regulators controlling the expression of genes needed for autotrophic growth are yet unknown. This information could benefit acetogen metabolic engineering and improve our understanding of their complex transcriptional regulation (Aklujkar et al., 2017; Marcellin et al., 2016; Nagarajan et al., 2013; Tan et al., 2013). Prediction of promoter motifs strictly based on computational analysis (based solely on the organism’s genome sequence) has the drawback of detection of promoter-like sequences across the genome, which is particularly pronounced in non-conserved DNA motifs (Patrik, 2006). An instrumental step towards more accurate promoter motif identification was the development of the differential RNA-sequencing (dRNA-Seq) technology, first described in 2010 by Sharma and colleagues (Sharma et al., 2010) for the human pathogen *Helicobacter pylori*.

dRNA-Seq enables the experimental determination of transcription start sites (TSSs) and correct mapping of TSSs enables genome-wide identification of promoters and gene expression regulatory sequences, besides providing experimental data for a more accurate genome annotation. Once a TSS has been experimentally determined, promoter sequences can be mapped from there. Thus, characterisation of the transcriptional architecture (i.e. TSSs and promoter motifs) and a more accurate annotation of acetogen genomes have the potential to yield valuable insights into the complex transcriptional regulation of acetogens. To date, only one study has determined TSSs in acetogens, using *Eubacterium limosum* (Song et al., 2017). Here, we used dRNA-Seq as a tool to identify the TSSs in the model-acetogen *Clostridium autoethanogenum* grown under autotrophic and heterotrophic conditions. The subsequent search for promoter motifs detected a previously undescribed motif associated with essential genes in acetogens. We then provide experimental evidence for the relevance of this new promoter motif (names hereafter P_cauto_) by identifying a TetR-family protein that activates gene expression from this motif by directly binding to the RNA polymerase.

## 2. Materials and Methods

### 2.1 Bacterial strains and growth conditions

*Clostridium autoethanogenum* strain DSM 10061 was obtained from The German Collection of Microorganisms and Cell Cultures (DSMZ). Cells were grown as described before (Marcellin et al., 2016). Briefly, heterotrophic and autotrophic growth were investigated in serum bottles on fructose (5 g/L) and on steel mill off-gas (35% CO, 10% CO_2_, 2% H_2_ and 53% N_2_), respectively. Cells were grown at 37 °C on a shaker (100 RPM, rounds per minute) and sampled for dRNA-Seq analysis from the exponential growth phase (OD_600nm_= 0.5-0.6).

### 2.2 Differential RNA-sequencing (dRNA-Seq)

Extraction and preparation of RNA for cDNA library construction were performed as described elsewhere (Marcellin et al., 2016). Briefly, RNA was extracted using TRIzol followed by column purification with RNAeasy (Qiagen). The resulting total RNA pools were sent to Vertis Biotechnologie AG (Freisig, Germany) for sequencing. The cDNA libraries were prepared using the 5′tagRACE method (Fouquier D’Hérouel et al., 2011). Firstly, the 5′ Illumina TruSeq sequencing adapter carrying sequence tag TCGACA was ligated to the 5′-monophosphate groups (5′P) of processed transcripts (TAP-on Figure 1A). Samples were then treated with Tobacco Acid Pyrophosphatase (TAP) to convert 5′-triphosphate (5′PPP) structures of primary transcripts into 5′P ends to which the 5′ Illumina TruSeq sequencing adapter carrying sequence tag GATCGA was ligated (TAP+ on Figure 1A). Next, first-strand cDNA was synthesised using an N6 randomised primer to which the 3′ Illumina TruSeq sequencing adapter was ligated after fragmentation.

**Figure 1.**
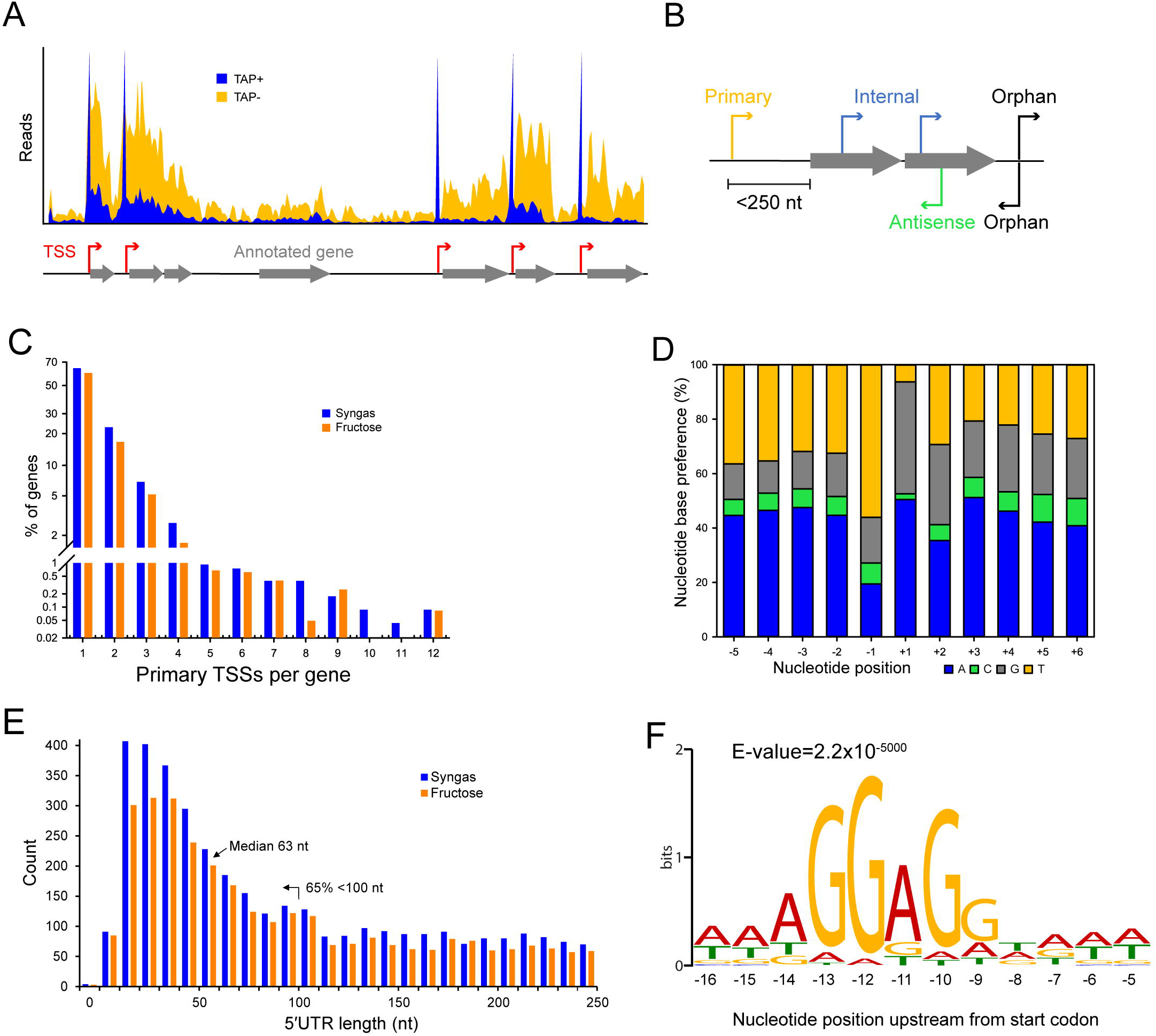
Characteristics of transcriptional and translational architecture in *C. autoethanogenum*. (A) Our dRNA-Seq approach generated genome-wide TSS maps through the comparison of libraries enriched for processed (TAP-) and primary (TAP+) transcripts. (B) Classification of TSSs for syngas and fructose as: primary, within 250 nt upstream of an annotated gene; internal, within an annotated gene; antisense, on the opposite strand of an annotated gene; orphan, not assigned to any of the previous classes. (C) Distribution of primary TSSs per gene for syngas and fructose. (D) Nucleotide base preference for transcription initiation from primary TSSs on syngas. +1 denotes the position of the TSS. (E) Distribution of 5′UTR lengths for primary TSSs for syngas and fructose. (F) The Shine-Dalgarno sequence AGGAGG is highly conserved within 9-14 nt upstream of the first start codon. Sequencing reads were processed with the TSSAR software (Amman et al., 2014) for automated *de novo* determination of TSSs from dRNA-Seq data using the following parameters: p-Value 1e-3, Noise threshold 10, Merge range 5. The Shine-Dalgarno sequence was searched 30 nt upstream of annotated genes (CP006763.1 and NC_022592.1) using the MEME software (Bailey et al., 2009) and the same parameters as for promoter motif search, except for-nmotifs 10, -maxw 30. See Methods for details.

The 5′ cDNA fragments were amplified with PCR using a proof reading enzyme and primers designed for TruSeq sequencing according to the manufacturer’s instructions. The main advantage of using the 5′tagRACE method (Fouquier D’Hérouel et al., 2011) for dRNA-Seq comes from amplifying the 5′ ends of processed and primary transcripts in a single PCR reaction, which preserves their quantitative representation in an RNA pool. Finally, 5′ cDNAs were purified using the Agencourt AMPure XP Kit (Beckman Coulter Genomics) and analysed by capillary electrophoresis before sequencing the single-end libraries using the Illumina NextSeq 500 system and a MID 150 Kit with 75 bp read length.

### 2.3 Determination of transcription start sites (TSSs)

Sequencing reads were aligned and mapped to the genome of *C. autoethanogenum* DSM 10061 (CP006763.1) using the software TopHat2 (Kim et al., 2013) without trimming or removal of any reads. Reads were processed with the TSSAR (*TSS A*nnotation *R*egime) software (Amman et al., 2014) for automated *de novo* determination of TSSs from dRNA-Seq data using the following parameters: p-Value 1e-3, Noise threshold 10, Merge range 5. The identified TSSs were classified as primary (within 250 nt upstream of an annotated gene), internal (within an annotated gene), antisense (on the opposite strand of an annotated gene), or orphan (not assigned to any of the previous classes) (Figure 1B). Since our main aim was the identification of the TSSs of essential genes for autotrophic growth in acetogens (e.g WLP), we focused on the primary TSSs.

### 2.4 Search for promoter motifs and the Shine-Dalgarno sequence

To determine promoter motifs, we searched for consensus sequence motifs 50 nt upstream of primary TSSs using the MEME software (Bailey et al., 2009) with the following parameters: – dna, -max size 10000000, -mod zoops, -nmotifs 50, -minw 4, -maxw 50, -revcomp, -oc. Only motifs with E-value ≤ 0.05 and at least 13 TSSs associated to it (i.e. at least two genes associated to it, Figure 1C) were considered and ranked based on the number of assigned TSSs (Supplementary file 1).

To search for the Shine-Dalgarno sequence, 30 nt upstream of annotated genes (CP006763.1 and NC_022592.1) were searched with the MEME software (Bailey et al., 2009) using the same parameters as in the promoter motif search, except for -nmotifs 10, -maxw 30.

### 2.5 Search for the new promoter motif in acetogens

Occurrence of the new promoter motif (see results) in *C. autoethanogenum*, *C. ljungdahlii*, *C. ragsdalei*, *C. coskatii*, *M. thermoacetica*, and *E. limosum* was determined using the FIMO tool (Grant et al., 2011) within the MEME software by searching for the sequence up to 300 nt upstream of annotated genes (since no TSS data is available) with default FIMO parameters. Occurrence in each acetogen relative to *C. autoethanogenum* was normalised with the number of annotated genes.

### 2.6 DNA-binding protein assay

Firstly, *C. autoethanogenum*—DSM 19630—cells were acquired from autotrophic bioreactor chemostat cultures (CO or CO+H_2_) described in a separate work (Valgepea et al., 2018). Briefly, cells were grown in bioreactor chemostat cultures in the chemically defined medium on either CO or CO+H_2_ at 37 °C, pH = 5, dilution rate of ~1 day^−1^ (µ~0.04 h^−1^), and at a biomass concentration ~1.4 gDCW/L. Cells were pelleted by immediate centrifugation (20,000 × *g* for 2 min at 4 °C), and stored at −80 °C until analysis.

Frozen pellets were thawed, resuspended in BS/THES buffer described in (Jutras et al., 2012) with pH adjusted to 7.0, and passed five times through the EmulsiFlex-C5 High Pressure Homogenizer (Avestin Inc.) according to the manufacturer’s instructions, with the final sample volume adjusted to 35 mL with the BS/THES buffer. Samples were then centrifuged (35,000 × *g* for 15 min at 4 °C) and the supernatant filtered using a 0.22 µM filter (Merck).

The DNA-binding protein assay was based on a pull-down/DNA affinity chromatography method described by Jutras and co-workers (Jutras et al., 2012) with the following modifications. The DNA sequences were of 125 bp length containing the respective promoter sequence in the middle with flanking regions downstream and upstream. pH of the buffers was adjusted to 7. The bait-target/ligand binding step was performed with 1 mL of cell extract without the addition of non-specific competitor DNA.

Next, either salmon sperm (Thermo) or Poly dI-dC (Sigma) were used as non-specific competitor DNA in the subsequent washing steps. Briefly, Dynabeads™ M-280 Streptavidin (Thermo Fisher Scientific) were mixed with DNA containing either the promoter sequence of CAETHG_1615, 1617 (WLP genes assigned with the new promoter motif), or 3224 (a glycolytic gene as a control for our assay since it was assigned the well-known TATAAT motif, which should yield binding of the RNAP and the housekeeping σ factor σ^A^). Next, the cell extract was added and samples were incubated for 30 min at room temperature. This was followed by two washing steps with the BS/THES buffer (Jutras et al., 2012) to remove proteins not bound to the target DNA. Finally, protein elution was performed in Tris-HCl (pH 7) with a successively increasing concentration of NaCl (200, 300, 500, 750 mM, 1M, and 2M). The eluted protein solutions were analysed by gel electrophoresis NuPAGE® Novex®Bis-Tris (Invitrogen) and visualized using Sypro® Ruby (Molecular Probes) according to the manufacturer’s instructions. The 500 mM NaCl eluate yielded the most prominent bands and therefore this eluate was used for further analysis. No bands were observed in the negative control when water was used instead of DNA (data not shown), confirming that the identified proteins were pulled down by the DNA sequences.

### 2.7 Protein digestion for mass spectrometry-based proteomics

For the digestion of proteins from gel band excision, the gel bands of interest were cut and de-stained for 1 h with a buffer of 50 mM ammonium bicarbonate (ABC) in 50% acetonitrile (ACN). Following buffer removal, 50 µL of 10 mM DTT was added and samples were incubated for 30 min at 60 ºC to reduce disulphide bonds. Next, the DTT solution was removed, and 50 µL of 55 mM iodoacetamide (IAA) was added and samples were incubated for 30 min in the dark at room temperature to alkylate sulfhydryl groups. After removal of the IAA solution, gel pieces were washed twice with 100 µL of 50 mM ABC, and dehydrated with 100% ACN. Protein digestion was performed overnight at 37 ºC by rehydrating gel pieces with 50 µL of Trypsin/Lys-C mix (10 ng/µL in 25 mM ABC) and 100 µL of ABC.

Extraction of peptides from gel pieces was performed by repeating the following steps five times: addition of 100 µL of 0.1% formic acid (FA) in 50% ACN and sonication of samples in a water bath for 10 min. Samples were then concentrated to near dryness using a centrifugal vacuum concentrator (Eppendorf) and resuspended in 50 µL of 0.1% FA. Finally, samples were desalted using C_18_ ZipTips (Merck Millipore) as follows: the column was wetted using 0.1% FA in 100% ACN, equilibrated with 0.1% FA in 70% ACN, and washed with 0.1% FA before loading the sample and washing again with 0.1% FA. Finally, peptides were eluted with 0.1% FA in 70% ACN, and then diluted 10-fold with 0.1% FA for mass spectrometry analysis.

For the digestion of proteins from the whole purified DNA bound material, the whole purified DNA-bound material from the DNA-protein binding assay was incubated for 30 min at 95 ºC. Next, 30 µL of 10 mM DTT was added and samples were incubated for 45 min at 55 ºC to reduce disulphide bonds. Then, 40 µL of 55 mM IAA was added and samples were incubated for 30 min in the dark at room temperature to alkylate sulfhydryl groups. Protein digestion was performed overnight at 37 ºC using 50 µL of Trypsin/Lys-C mix (10 ng/µL in 25 mM ABC) and stopped by lowering the pH to 3 using FA. Finally, the samples were desalted and prepared for mass spectrometry analysis as described above.

### 2.8 Protein identification using mass spectrometry

Detection of proteins in both the digestion products of gel band excision and the whole captured material was performed using a QTOF Sciex 5600 or a Thermo Orbitrap Elite mass spectrometer (depending on instrument availability) with details described elsewhere (Kappler and Nouwens, 2013) (Yang et al 2016) with a modified liquid chromatographic (LC) gradient. Protein identification was performed using the software ProteinPilot v5.0 (ABSciex) with the Paragon Algorithm against the NC_022592.1 and CP006763 genome annotations with the following search parameters: Trypsin+LysC digestion; IAA as cysteine alkylation; Thorough search effort; FDR analysis. Only proteins below 1% false discovery rate (FDR; estimated global) and with at least two peptides with more than 95% confidence were considered as identified.

### 2.9 Molecular Biology Techniques

The full list of bacterial strains, plasmids, and primers used in this work for the *in vivo* transcription assay and protein overexpression step are shown in Supplementary file 2. Luria-Bertani (LB) broth or agar with antibiotics were used for growth.

*E. coli* DH5α was used as the cloning strain and performed transformations according to the manufacturer’s instructions (BIOLINE). *E. coli* BL21 was used in the *in vivo* transcription assay and protein overexpression step. *E. coli* BL21 chemically competent cells were prepared using the RuCl_2_ method (Green and Rogers, 2013).

PCR amplification of targeted sequences was performed using the Phusion polymerase (NEB) and the OneTaq polymerase (NEB). Plasmid were assembled using standard ligation with the T4 DNA ligase or using Gibson assembly (Gibson et al., 2009).

#### 2.9.1 Construction of a σ-factor candidate expression system in *E. coli*

Candidates for potential σ factors were selected based on protein identification using mass spectrometry (see above) from proteins annotated as transcriptional regulatos (Table 1). Additionally, we also built a plasmid for the L-seryl-tRNA(Sec) selenium transferase (CAETHG_2839) (identified as a stronger band in the pull-down assay (Figure 3B)), and the housekeeping σ in *Clostridia* (σ^A^) (CAETHG_2917) (Figure 3B).

**Table 1.** *C. autoethanogenum* proteins annotated as transcriptional regulators uniquely binding to the new promoter motif P_cauto_

| Gene ID<sup>a</sup> | Gene ID<sup>b</sup> | Gene product annotation<sup>a</sup> |
| --- | --- | --- |
| CAETHG_RS02185 | CAETHG_0459 | TetR/AcrR family transcriptional regulator |
| CAETHG_RS04465 | CAETHG_0936 | TetR/AcrR family transcriptional regulator |
| CAETHG_RS19205 | CAETHG_3915 | GntR family transcriptional regulator |

<sup>a</sup> Annotation NC\_022592.1.
<sup>b</sup> Annotation CP006763.1 (Brown et al., 2014).

The potential σ factor candidates were cloned into plasmid pET28a+ to be expressed under the control of a T7 promoter. DNA sequences were PCR amplified using the primers shown in Supplementary file 2 and purified using a QIAGEN kit. Next, the plasmid pET28a+ was linearised using restriction enzymes NdeI and HindIII and purified using a QIAGEN kit. Codon optimisation was required to express the σ factor candidates of TetR-family protein (CAETHG_0459) and σ^A^ (CAETHG_2917) before DNA sequences were synthesised as gene block (gBlock®) fragments.

Plasmids with the σ factor candidates were then assembled by Gibson assembly using equimolar concentrations of the linearised backbone plasmid and the PCR fragment in a 20 µL reaction. After incubation at 50 ºC, 5 µL of the Gibson mix was then used to transform *E. coli* DH5α by heat shock. After recovery on SOC media at 37º C for 60 min, 100 µL of cells were spread on LB agar plates containing kanamycin (50 µg/mL). Plates were then incubated at 37 ºC for 16 h and kanamycin resistant colonies were tested by colony PCR for proper assembly using pET_conf(FWD)/pET_conf(REV) primers (Supplementary file 2). A colony that tested positive for assembly was then picked and grown overnight on LB media containing kanamycin. Plasmids were recovered from 5 mL of overnight culture using a QIAGEN miniprep kit and the digestion profile was verified with the assembly. Plasmids were then used to transform *E. coli* BL21 chemically competent cells (described above).

*E. coli* BL21 strains harbouring σ factor candidate-expressing plasmids were then grown overnight on LB media containing kanamycin and 1 mM IPTG (Isopropyl β-D-1-thiogalactopyranoside). Next, 2 mL of overnight culture were spun down and the supernatant was removed. Next, the cell pellet was resuspended in the BugBuster master mix solution (Novagen) for protein extraction following the manufacturer’s instructions. The insoluble and soluble fractions were loaded into an SDS-PAGE gel to confirm the overexpression of the σ factor candidates (data not shown).

#### 2.9.2 Construction of a P_cauto__GFP-UV reporter fusion system in *E. coli*

To determine whether the σ factor candidates could activate transcription, we assembled a GFP-based reporter expression system under the control of the P_cauto_. Firstly, plasmid pBR322 was digested with HindIII, purified, and used as the backbone followed by PCR amplification of the DNA sequence containing P_cauto_ from the *C. autoethanogenum* genome (500 bp upstream of the start codon of the gene CAETHG_1617) and purification using a QIAGEN kit.

Next, the GFP-UV gene was PCR amplified from plasmid pBR_PprpR-GFPUV and purified after which the three DNA fragments were added at an equimolar concentration to a Gibson assembly mix subsequently incubated at 50 ºC. 5 µL of the Gibson mix weas used to transform chemically competent *E. coli* DH5α cells by heat shock and after incubation at 37 ºC, 100 µL of cells were spread on LB agar plates containing ampicillin (100 ug/mL) and incubated at 37 ºC for 16 h. Ampicillin resistant colonies were then tested by colony PCR using the primer sets of P_cauto_-GFP_conf(FWD-1)/ P_cauto_-GFP_conf(REV-1) and P_cauto_-GFP_conf(FWD-2)/ P_cauto_-GFP_conf(REV-2) (supplementary file 2). Confirmed colonies were picked and grown overnight on LB containing antibiotic for plasmid recovery. The digestion profile confirmed the assembly of plasmid pBR_P_cauto__GFP.

The P_cauto_-GFP-UV was excised from pBR_P_cauto__GFP using restriction enzyme HindIII. Digestion mix was loaded on a 1% agarose gel and the P_cauto_-GFP-UV region recovered using a QIAGEN gel extraction kit. Then, the recovered DNA sequence was cloned into plasmid pACYC184, which was previously digested with HindIII and purified using a QIAGEN kit.

Ligation was performed according to the manufacturer’s instruction and 5 µL of the mix was used to transform *E. coli* DH5α competent cells. After heat shock and incubation, 100 µL of cells were spread on LB agar containing chloramphenicol (30 ug/mL) and incubated at 37 ºC for 16 h. Chloramphenicol-resistant colonies were tested by colony PCR for proper assembly. Positive colonies were then grown overnight on LB media and the plasmid was recovered. Assembly of plasmid pACYC_P_cauto__GFP was confirmed by digestion profile and Sanger sequencing (AGRF, Australia) (data not shown).

#### 2.9.3 Construction of a variants for the P_cauto_ promoter motif region

Later a new reporter system including the P_cauto_ and the GFPuv sequences was built to remove the 500 bp upstream region in pBR_P_cauto__gfp. The idea was to keep only the sequence used for the pull-down assay plus including the ribosomal binding site (Shine-Dalgarno sequence) to be tested *in vivo* with TetR-family protein (CAETHG_0459) and σ^A^ (CAETHG_2917) (see net section), the two proteins that responded positively in the *in vivo* assay (see results). This new plasmid, pBR_P_cauto_130_gfp, was built by cloning the PCR product of primers WLP130F and WLP130R using pBR_P_cauto__gfp as template, at the HindIII site of pBR322 by Gibson assembly (supplementary file 2). Then, the P_cauto_130_gfp region was excised from pBR_P_cauto_130_gfp using HindIII and ClaI, and cloned by ligation in pACYC184 to build plasmid pAC_P_cauto_130_gfp. A variation of the promoter region (pAC_P_cauto_30C_gfp) was also built to introduce single nucleotide changes in the WLP promoter motif. Changes were as follow: ctggagcaggttttgtagttgcagtaactggttcaata, changed to ccatcaaaggtcttaaagttgcagtaactggttcaata. This promoter was again tested with the TetR-family protein (CAETHG_0459) and σ^A^ (CAETHG_2917). All plasmids maps used can be found in supplemental information.

#### 2.9.4 *In vivo* transcription activation of Pcauto-GFP(UV) fusion by the candidate genes in *E. coli*

*E. coli* BL21 was used for the *in vivo* assay. Firstly, six biological replicate cultures of cells were grown in a 96-well plate (Corning Costar catalogue number #3799) carrying the pACYC plasmid with or without (to correct for the autofluorescence of the cells) the promoter-GFPuV fusion reporter in *trans* with a pET plasmid carrying each of the σ factor candidates. Additionally, a system with cells carrying either the pACYC promoter-GFPuV fusion reporter or its backbone plasmid plus the pET plasmid with no candidate was used as the control.

Cells were grown in 150 µL of LB media containing kanamycin and chloramphenicol at 30 ºC and agitation of 200 RPM. At mid-exponential phase, cells were sub-cultured to a black 96-well plate (Greiner #655090) to an initial OD of 0.05-0.1 in LB media containing kanamycin and chloramphenicol supplemented with either 0.0 mM IPTG (No IPTG) or 1.0 mM IPTG. The *in vivo* experiment was performed at 30 ºC and agitation of 200 RPM.

Growth was followed by measuring the optical density (OD) at 600 nm while fluorescence intensity (FI; for GFP expression) was measured using the excitation filter of 355 nm and an emission filter of 520 nm. The experiment was conducted using the FLUOstar Omega microplate reader and the Omega software v.1.20 (BMG LabTech). Fluorescence intensity was normalized per OD (FI/OD) and the signal resulting from the cells harbouring the backbone plasmid only was subtracted from the cells carrying the promoter-GFP fusion reporter (Normalized FI/OD).

For the WLP promoter motif variants (described in the previous sentence) four biological replicates were used.

Student’s t-test (two-tailed) was performed between each of the candidate’s normalized FI/OD value without and with IPTG and between the control system. A candidate gene was considered to activate gene expression from P_cauto_ if it met both of the following two conditions: 1) there was a statistically significant difference (p-value<0.01) in FI/OD between the candidate without and with IPTG; 2. there was a statistically significant difference (p-value<0.01) between the FI/OD signal of the candidate and the control vector (PET_) with IPTG.

#### 2.9.5 Overproduction and purification of TetR-family protein (CAETHG_0459)

To enable the test whether the TetR-family protein CAETHG_0459 activates transcription from P_cauto_ by interacting directly with the RNAP, the target protein had to be heterologously expressed and purified for the protein-protein interaction assay (see 2.9.5).

The *E. coli* strain harbouring the plasmid pET_TetR1 (CAETHG_0459) was grown at 30 º C and 200 RPM until mid-exponential phase in LB media containing kanamycin. Cells were sub-cultured to 1 L LB media containing kanamycin to an initial OD of 0.05-0.1 and subsequently grown until OD ~1 at 30 ºC and 200 RPM. Then, 1.0 mM IPTG was added and cells were left growing until OD ~3. Cells were pelleted from 1 L culture by centrifugation at 5,000 × *g* for 20 min at 4 °C, the pellet was resuspended in 5 mL of the BugBuster Master Mix (Merck Millipore #71456) per gram of wet cell weight with EDTA-free protease inhibitor cocktail (Sigma #11836170001), and then incubated in a rotating mixer for 20 min at room temperature. Next, cells debris were removed by centrifugation at 16,000 × *g* for 20 min at 4 °C and the supernatant (supplemented with 20 mM Imidazole) was loaded on a 1 mL Ni^+^-HisTrapHP column (GE Healthcare #71-5027-68 AK) and washed with a buffer containing 100 mM Tris-HCl (pH 7), 100 mM NaCl, 20 mM Imidazole.

The TetR-family protein protein CAETHG_0459 was eluted in the same wash buffer containing a stepwise imidazole gradient (50-500 mM) following a buffer exchange performed using a HiTrap Desalting column (GE Healthcare #17-1408-01). Finally, the purified protein was stored in 50 mM Na_2_HPO_4_, 300 mM NaCl, pH7, 50% glycerol. Protein purity was analysed by gel electrophoresis using NuPAGE® Novex®Bis-Tris (Invitrogen) and stained with SimplyBlue™ SafeStain (Novex). Protein concentration was measured by the Direct Detect Spectrometer (Merck Millipore).

#### 2.9.6 TetR-family protein (CAETHG_0459)-RNA polymerase Core enzyme interaction experiment

The protein-protein interaction (PPI) experiment was performed as described previously (Raffestin et al., 2005) with some modifications. The purified TetR-family protein (CAETHG_0459) with 6-His-tag (2 µg) was coupled to Ni+-NTA agarose beads (Thermo #88831) in 800 µL of buffer A (50 mM Na_2_HPO_4_, 300 mM NaCl, 50 mM imidazole, pH 7). The beads coupled with the taret protein were then washed three times in buffer B (50 mM Na_2_HPO_4_, 300 mM NaCl, 0.1% Tween 20, 50 mM imidazole, pH 7). Next, the beads-protein complex was incubated with *E. coli* RNA polymerase Core enzyme (2.5 µg) (BioLabs #M0550S) at 37 ºC for 2 h. After two washes in buffer A, the beads-protein complex was suspended in 15 µL of Laemmli Buffer (32.9 mM Tris-HCl, pH6.8, 13.15 % (w/v) glycerol, 1.05 % SDS, 0.005% bromophenol blue, 355 mM 2-mercaptoethanol), heated at 100 ºC for 5 min, and analysed by gel electrophoresis using NuPAGE® Novex®Bis-Tris (Invitrogen) and stained with SimplyBlue™ SafeStain (Novex). The negative control was performed by incubating the RNA polymerase Core enzyme with Ni+-NTA agarose beads following the same procedure.

#### 2.9.7 Visualization of cells harbouring the Pcauto-GFP(UV) fusion and the TetR-family protein (CAETHG_0459) plasmids by microscopy

Cells carrying the Pcauto-GFP(UV) fusion reporter and the TetR-family protein (CAETHG_0459) plasmids were analysed by microscopy to visualize the expression of GFP. For this, cells were plated in an LB agar plate (LB media containing 6 g/L of agar) containing 1.0 mM IPTG, kanamycin, and chloramphenicol. After overnight incubation at 37º C, colonies were visualized using the ZOE™ Fluorescent Cell Imager (Bio-Rad) using the manufacturer’s instructions and following parameters: Gain: 40; Exposure (ms): 340; LED intensity: 22; Contrast: 59.

## 3. Results

### 3.1 Differential RNA-sequencing (dRNA-Seq)

In this work, we aimed to determine the TSSs of essential genes for autotrophic growth of the model-acetogen *C. autoethanogenum* (e.g. genes in the WLP and of hydrogenases). We thus performed dRNA-Seq analysis (Sharma et al., 2010) of autotrophic (CO, CO_2_, and H_2_; referred to as ‘syngas’) and heterotrophic (fructose) cultures of *C. autoethanogenum* to experimentally determine TSSs and promoter motif(s) associated with essential genes for autotrophic growth in acetogens.

Previously described batch cultures (Marcellin et al., 2016) were sampled during exponential growth and subjected to dRNA-Seq cDNA library preparation and sequencing. The cDNA libraries were prepared using the 5′tagRACE method (Fouquier D’Hérouel et al., 2011), an improved library preparation method compared to TEX (5’-phosphate-dependent Terminator RNA exonuclease) that has the advantage of preserving the quantitative representation of 5′ ends between processed (5’-P end) and primary (5’-PPP end) transcripts (see Methods). TSSs were determined by comparing the libraries enriched for processed (TAP-) and primary (TAP+) transcripts (Figure 1A) using the TSSAR tool (Amman et al., 2014).

### 3.2 Overall dRNA-Seq features of *C. autoethanogenum*

We classified TSSs as primary, internal, antisense, and orphan (Figure 1B, Table S1) and found primary TSSs only for around half of the annotated genes (3,983) in *C. autoethanogenum* (Brown et al., 2014) (Table S1). More than 60% of the genes contain only one primary TSS, while the rest show up to 12 TSSs (Figure 1C, Table S1). Focusing on the 14 main metabolic groups of *C. autoethanogenum* genes as described in (Brown et al., 2014), we detected primary TSSs for all genes except for the Nfn transhydrogenase complex (CAETHG_1580) (Table S2). While primary TSSs were detected for seven of the 11 genes of the WLP biosynthetic gene cluster (CAETHG_1606-21), only half of the WLP TSSs were shared between syngas and fructose. For example, genes of the WLP methyl branch (CAETHG_1614-17) contained 20 primary TSSs on syngas compared to only nine on fructose. On the other hand, the TSSs associated with Hydrogenases and ATPase genes were found in similar numbers between syngas and fructose.

Determination of nucleotide base preferences for transcription initiation within five nucleotides downstream and upstream of the primary TSSs showed a clear enrichment of adenine (A) and guanine (G) at +1 (~90%) and thymine (T) at −1 for both syngas (Figure 1D) and fructose (data not shown). Overall, adenine and cytosine were the most and least preferred nucleotide bases, respectively.

Analysis of 5′untranslated regions (5′UTRs)—the sequence between the TSS and the annotated start codon—indicates transcripts potentially associated with post-transcriptional regulation and thus of mRNA stability and translational efficiency (Cho et al., 2014). Calculation of 5′UTR lengths for primary TSSs showed a median length of 63 nt with 65% of TSSs <100 nt for both growth conditions (Figure 1E and Table S1). Genes with longer UTR lengths tend to be regulated more at the post-transcriptional level (Cho et al., 2009; David et al., 2006). On the other hand, leaderless mRNAs—mRNAs with no or <10 nt 5′UTR—are translated in the absence of upstream signals (typically the Shine-Dalgarno sequence) (Shine and Dalgarno, 1974; Zheng et al., 2011) used for regulating translational efficiency through ribosome binding. We found ~70 (~2%) leaderless mRNAs with <10 nt 5′UTRs, none of which were in the WLP, Hydrogenases, Acetate or Ethanol groups (Figure 1E and Table S3).

In addition to the ability to determine TSSs, dRNA-Seq analysis also facilitates a more accurate annotation of the genome. Based on the TSSs and the Shine-Dalgarno (AGGAGG) position that was found to be highly conserved within 9-14 nt upstream of the first start codon (ATG/CTG/GTG/TTG) (Figure 1F), we re-annotated the start codon for 38 genes and confirmed the changes in one gene by peptide identification using mass spectrometry (Table S4). Moreover, either the start or stop codon of an additional 99 genes, which had previously been annotated in different frames, were manually corrected. The corrections have been deposited into NCBI under the accession number BK010482 and the complete manually corrected genbank file of *C. autoethanogenum* is available in Table S5.

### 3.3 Discovery of a new promoter motif

The RNA polymerase (RNAP) needs to form a holoenzyme with a σ factor in bacteria to recognise a specific promoter motif (sequence) and initiate transcription (Feklistov et al., 2014; Gruber and Gross, 2003). Experimentally determined TSS data from dRNA-Seq analysis is ideal for *in silico* determination of promoter motifs, which is important for understanding transcriptional regulation, especially in less-studied bacteria such as acetogens.

We searched for consensus sequence motifs 50 nt upstream of primary TSSs using the MEME software (Bailey et al., 2009) and were able to determine seven promoter motifs in *C. autoethanogenum* (E-value ≤ 0.05) (Tables S6 and 7 for syngas and fructose growth, respectively). Of those identified, only three motifs were assigned with more than 100 TSSs and shared between the two datasets, likely representing the most conserved motifs in *C. autoethanogenum* (Figure 2A).

**Figure 2.**
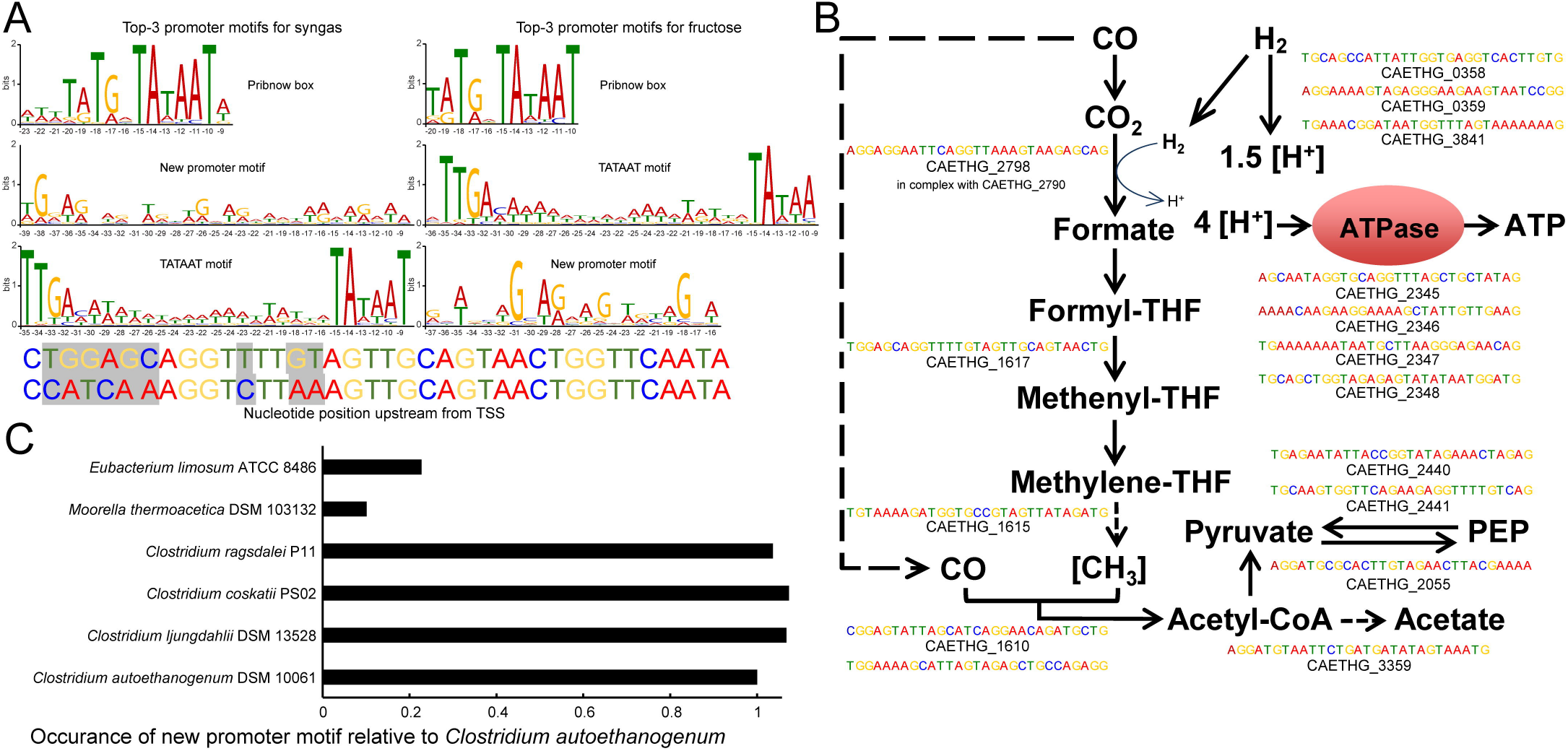
*In silico* determination of genome-wide promoter motifs in *C. autoethanogenum*. (A) The top-3 promoter motifs for primary TSSs are shared among syngas and fructose. The height of the letter indicates its relative frequency at the given position within the motif. Refer to Tables S5-8 for all the determined motifs and their assigned TSSs. The mutated nucleotides used in the *in vivo* assay for P_cauto_ motif are also shown. We show the nucleotide position relative to the TSS in all top3 motifs (B) The new promoter motif (P_cauto_) is assigned with TSSs of essential genes in acetogens. Motifs with the lowest p-value for syngas are shown. Refer to Tables S2 and 5-8 for all the TSSs and genes associated with P_cauto_. (C) The P_cauto_ motif is represented in other industrially relevant acetogens. Occurrence in each acetogen relative to *C. autoethanogenum* is normalised with the number of annotated genes. To determine promoter motifs in *C. autoethanogenum*, we searched for consensus sequence motifs 50 nt upstream of primary TSSs using the MEME software (Bailey et al., 2009) with the following parameters: -dna, -max size 10000000, -mod zoops, -nmotifs 50, -minw 4, -maxw 50, -revcomp, -oc.

The top motif was found 10 nt upstream of primary TSSs (447 and 543 TSSs for syngas and fructose, respectively; E-value<10^−111^) and resembles the Pribnow box (TATGnTATAAT), which is associated with the housekeeping σ factors of *Escherichia coli* (σ^70^; (Walker and Osuna, 2002)), *Helycobacter pylori* (σ^80^; (Sharma et al., 2010)) and *Clostridium acetobutylicum* (σ^A^; (Sauer et al., 1994, 1995)). Expectedly, the well-known −35 TTGACA and −10 TATAAT motifs (TATA box in eukaryotes and archaea) for housekeeping σ factors (Burgess and Anthony, 2001) was also among the top-3 promoter consensus sequences (392 and 262 TSSs for syngas and fructose, respectively; E-value<10^−46^). These two motifs were assigned for most of the genes of glycolysis/gluconeogenesis and the TCA cycle (Table S2).

The third most abundant promoter motif has, to the best of our knowledge, not previously been reported in the literature (Figure 2A). P_cauto_, is highly conserved both during growth on syngas (Motif 02 in Table S5; 392 TSSs; E-value<10^−174^) and fructose (Motif 03 in Table S6; 224 TSSs; E-value<10^−77^). Importantly, P_cauto_ seems to be involved in the transcriptional regulation of essential genes for acetogens and was assigned to genes of the WLP cluster (CAETHG_1606-21) and the metabolic groups, as described in (Brown et al., 2014), of Hydrogenases, Acetate, ATPase, and Pyruvate (Figure 2B; Tables S2, S6, and S7). We confirmed the unique presence of the “new promoter motif” upstream of the TSSs. Investigation of its upstream regions up to 100 or 150 nt showed no other motif apart from the one conserved within 50 nt upstream of TSSs. This new promoter is well characterised by an evenly interspaced (A/T)G repetition with an almost central A/T position (Figure S1). These observations potentially indicate the presence of a new σ factor or transcriptional regulator of critical importance in acetogens.

### 3.4 RNA polymerase and proteins annotated as transcriptional regulators specifically bind P_cauto_

We performed DNA-protein binding assays to determine if the RNAP and/or other protein(s) bind to P_cauto_. The promoter sequences of two WLP genes (CAETHG_1615 and 1617, Methylene-tetrahydrofolate reductase domain-containing protein and Methenyltetrahydrofolate cyclohydrolase, respectively) annotated with P_cauto_ were used for the DNA-protein binding assay using the promoter pull down/DNA affinity chromatography method (Figure 3A; (Jutras et al., 2012)). The promoter sequence of a glycolytic gene (CAETHG_3424, glyceraldehyde-3-phosphate dehydrogenase, type I) was included as a control for the assay since it was assigned the well-known TATAAT motif, which should yield binding of the RNAP and the housekeeping σ factor, σ^A^. DNA-bound proteins captured using streptavidin-coupled magnetic Dynabeads™ were identified using mass spectrometry of the digestion products of the whole captured material and of gel band excisions. Since this DNA-protein binding assay requires significant amounts of cellular protein material, especially for efforts to identify low abundance proteins such as σ factors or transcriptional regulators, autotrophic bioreactor chemostat cultures (CO or CO+H_2_) of *C. autoethanogenum* described in a separate work (Valgepea et al., 2018) were sampled for this analysis.

**Figure 3.**
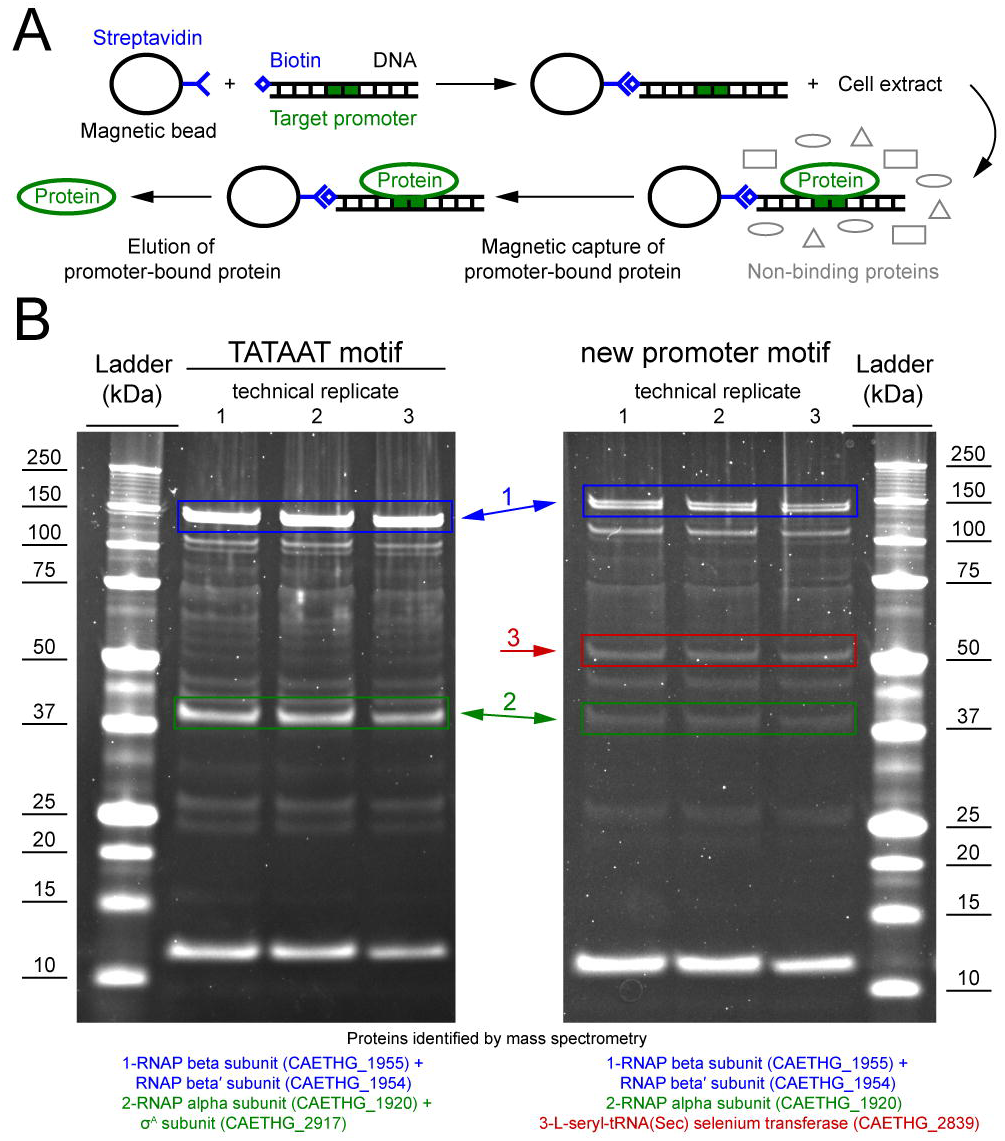
DNA-protein binding assay shows specific binding of *C. autoethanogenum* RNAP subunits and a selenium transferase to the new promoter motif. (A) Overview of the DNA-protein binding assay (i.e. the promoter pull down/DNA affinity chromatography method (Jutras et al., 2012)). (B) Separation of proteins specifically bound to the TATAAT motif (for gene CAETHG_3424) or the new promoter motif (for gene CAETHG_1617) with gel electrophoresis and identification using mass spectrometry. The alpha and beta subunits of the RNAP (CAETHG_1920 and 1954-55) were successfully identified for both the new promoter motif (CAETHG_1615 and CAETHG_1617) and the TATAAT motif control. Technical replicate denotes replicate of the DNA-protein binding assay (panel A) (data not shown for CAETHG_1615).

The promoter pull down/DNA affinity chromatography method (Figure 3A; (Jutras et al., 2012)) was fine-tuned for *C. autoethanogenum*. Eluting the proteins with 500 mM NaCl yielded the most prominent bands while no bands were observed in the negative control when water was used instead of DNA (data not shown), which confirms that the identified proteins were pulled down by the DNA sequences (see Methods). The alpha and beta subunits of the RNAP (CAETHG_1920 and 1954-55) were successfully identified for both P_cauto_ (CAETHG_1615 and CAETHG_1617) and the TATAAT motif control (Figure 3B). Additionally, the RNAP omega subunit was identified in the whole purified DNA-bound material for both motifs (Table S8). The housekeeping σ^A^ (CAETHG_2917) was detected for the TATAAT motif control as expected (Figure 3B). A stronger band was identified in the P_cauto_ gels around 50 kDa and identified as a protein annotated as L-seryl-tRNA(Sec) selenium transferase (CAETHG_2839; 51.5 kDa) (Figure 3B). Finally, mass spectrometry analysis of the whole purified DNA-bound material identified three proteins annotated as transcriptional regulators (based on NC_022592.1) that were unique for the P_cauto_ (Table 1) and found for both CO and CO+H_2_ cultures across technical replicates of the DNA-protein binding assay (Table S8).

### 3.5 TetR-family transcriptional regulator (CAETHG_0459) activates transcription from P_cauto_ *in vivo*

To determine whether any of the three identified protein candidates annotated as transcriptional regulators that uniquely bind to P_cauto_ (Table 1) could activate transcription from this promoter, we created a transcriptional fusion reporter vector harbouring the sequence of P_cauto_ in-frame with a green fluorescence protein (GFPuV). We also tested transcriptional activation using the L-seryl-tRNA(Sec) selenium transferase (CAETHG_2839) (identified as a stronger band in the pull-down assay (Figure 3B)), and using the housekeeping σ factor in clostridia (σ^A^) (CAETHG_2917), since it has been reported that promoter binding sites of different σ factors can overlap (Cho et al., 2014). Transcriptional activation of P_cauto_ with concomitant GFP production was investigated in *E. coli* by inducing the expression of the candidate activator proteins from a second T7 protein over-expression vector cloned into plasmid pET28e+ by the addition of IPTG (see Methods). Fluorescence was measured at early-exponential growth (OD ~0.26) as FI/OD.

After subtracting the signal from cells harbouring the two plasmids but lacking the fusion reporter (promoter + GFP, see Methods), only induction of the TetR-family transcriptional regulator protein (CAETHG_0459) (out of the three transcriptional regulator candidates) led to statistically higher levels of GFP expression (p<0.01) compared to the control vector with no candidate (Figure 4A and Table S9). Interestingly, induction of σ^A^ also led to transcription activation (p<0.01). We then confirmed expression of GFP in the strain expressing CAETHG_0459 grown on a plate with IPTG using fluorescence microscopy (Figure 4B). This shows that both CAETHG_0459 and σ^A^ independently activate transcription from P_cauto_. Importantly, the motif is associated with the expression of essential genes in gas-fermenting acetogens including genes in the WLP and hydrogenases (Table S2, S6-7).

**Figure 4.**
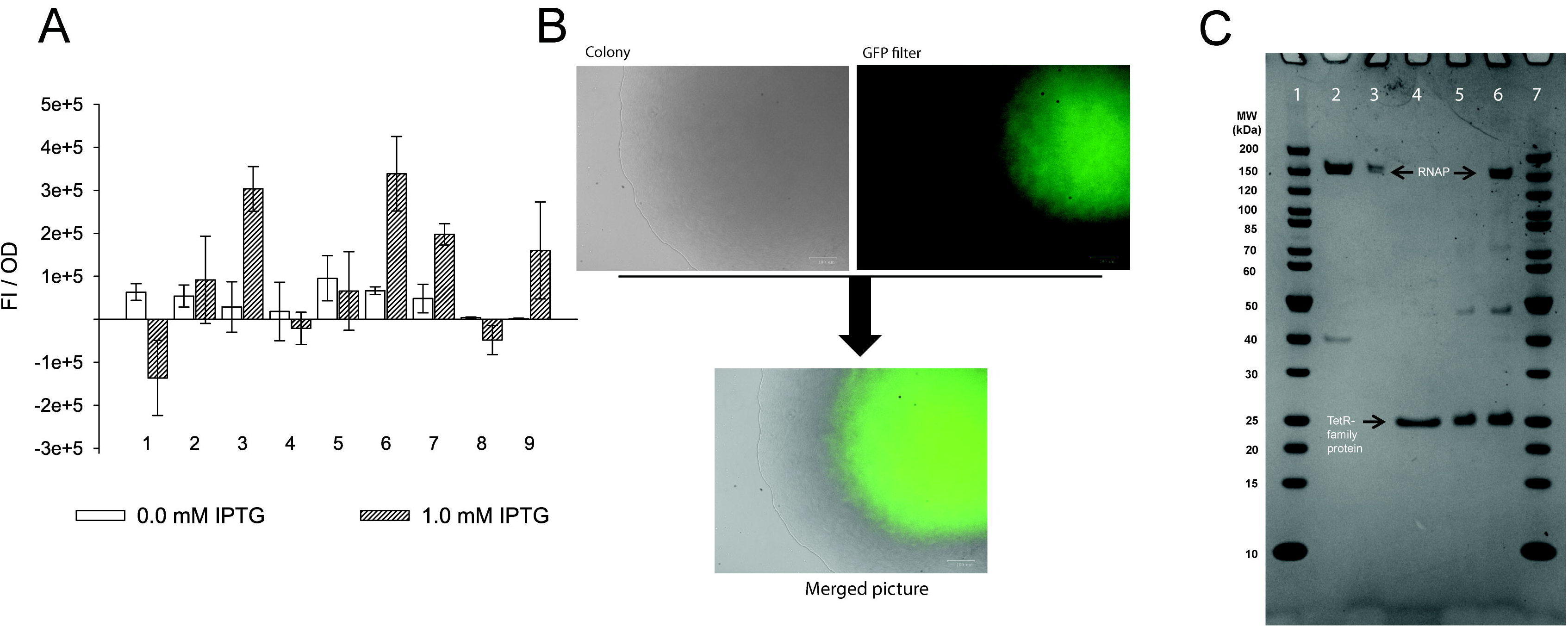
TetR-family transcriptional regulator (CAETHG_0459) and σ^A^ (CAETHG_2917) activate expression from the new promoter motif. (A) *In vivo* experiment using *E. coli* cells carrying the pACYC plasmid with the new promoter-GFPuV fusion report in *trans* with a pET plasmid carrying each of the candidates. The experiment was conducted with either 0.0 mM or 1.0 mM IPTG. Only in the presence of TetR-family protein (CAETHG_0459) and σ^A^ (CAETHG_2917) the fluorescence intensity normalized per OD (FI/OD) is statistically significantly different (p-value <0.01) compared to the control system (with no candidate protein). 1 Cells harbouring the PET_ (Negative control with no candidate gene); 2 Selenium transferase (CAETHG_2839); 3 TetR-family protein (CAETHG_0459); 4 TetR-family protein (CAETHG_0936); 5 GntR (CAETHG_3915); 6 σ^A^ (CAETHG_2917); 7 Short version (130 bp) of pAC_P_cauto_30C_gfp and TetR-family protein (CAETHG_0459); 8 Mutated version of the promoter region (pAC_P_cauto_30C_gfp) by introducing nucleotide changes as follow: ctggagcaggttttgtagttgcagtaactggttcaata, changed to ccatcaaaggtcttaaagttgcagtaactggttcaata and TetR-family protein (CAETHG_0459); 9 Short version (130 bp) of pAC_P_cauto_30C_gfp and σ^A^ (B). Cells carrying the TetR-family protein (CAETHG_0459) grown in LB-agar plate with 1 mM IPTG were visualized under microscopy for fluorescence (GFP) visualization. (C) Protein-protein interaction assay. TetR-family protein (CAETHG_0459) was incubated with *E. coli* RNA polymerase Core enzyme. Lane 1: Marker (Thermo #26614); Lane 2: *E. coli* RNA polymerase Core Enzyme; Lane 3: *E. coli* RNA polymerase Core incubated with Ni+ agarose beads and washed; Lane 4: Purified TetR-family protein (CAETHG_0459); Lane 5: Ni+ agarose beads coupled with TetR-family protein (CAETHG_0459); Lane 6: Ni+ agarose beads coupled with TetR-family protein (CAETHG_0459) incubated with RNA polymerase Core and washed; Lane 7: Marker

The 130bp variant (which includes the sequence used for the pull-down assay plus the ribosomal binding site) also showed statistically significance (p-value <0.01) of fluorescence increase when TetR-family transcriptional regulator protein (CAETHG_0459) was present. Similarly, σ^A^ could also activate transcription, however only at the level of p-value <0.05. Interestingly when mutations were included in the promoter motif, TetR-(CAETHG_0459) could no longer activate expression of GFP, as expected (Figure 4A).

### 3.6 CAETHG_0459 directly binds to the RNA polymerase core enzyme

As TetR-family proteins often act as transcriptional regulators (Cuthbertson and Nodwell, 2013), we next investigated whether TetR-family protein CAETHG_0459 activates transcription from P_cauto_ by interacting directly with the RNAP. Transcriptional regulators can reversibly interact with the RNAP Core enzyme independently of a DNA sequence to help activate transcription from a range of promoters (Burgess and Anthony, 2001; Feklistov et al., 2014). We thus performed an *in vitro* protein-protein interaction assay to test whether protein CAETHG_0459 directly interacts with RNA polymerase Core in the absence of DNA. The purified His-tagged CAETHG_0459 protein linked to Ni^2+^-beads was incubated with the RNAP Core enzyme (see Methods). SDS-PAGE analysis clearly demonstrated an interaction between the core RNA polymerase and CAETHG_0459 (Figure 4C lane 6) and shows that CAETHG_0459 acts as a positive transcriptional regulator that activates transcription from P_cauto_ by directly binding to the RNAP.

### 3.7 P_cauto_ is represented in other acetogens

We next investigated if P_cauto_ was represented in other industrially relevant acetogens with available genomes: *Clostridium ljungdahlii*, *C. ragsdalei*, *C. coskatii*, *Moorella thermoacetica*, and *Eubacterium limosum* (Bengelsdorf et al., 2016; Redl et al., 2017; Shin et al., 2016; Song et al., 2017). We performed the reverse of the methodology previously used to search for consensus sequence motifs by looking for the occurrence of P_cauto_ 300 nt upstream of annotated genes (since no TSS data was available) using the FIMO tool (Grant et al., 2011) within MEME. As expected based on their phylogenetic proximity (Bengelsdorf et al., 2013; Brown et al., 2014; Shin et al., 2016), *C. ljungdahlii*, *C. ragsdalei*, and *C. coskatii* showed similar occurrences of P_cauto_ (Figure 2C). Interestingly, while the representation in *M. thermoacetica* was very low, P_cauto_ seems to be present also in *E. limosum*. This result highlights the need for experimental determination of TSSs in more acetogens.

## 4. Discussion

Acetogens offer an enormous potential for the production of fuels and chemicals from gaseous waste feedstocks (Claassens et al., 2016; Dürre and Eikmanns, 2015; Liew et al., 2016; Molitor et al., 2016), with ethanol already being produced at industrial scale by LanzaTech. Acetogens have two major carbon fixation pathways: the WLP for autotrophic growth and glycolysis for heterotrophic growth. Although both the WLP and glycolysis/gluconeogenesis pathways operate during autotrophic and heterotrophic growth, the WLP carries a substantially higher metabolic flux during autotrophy (Valgepea et al., 2017a, 2018) and *vice versa* (Valgepea et al., 2017b). We and others have shown that transcriptional regulation between autotrophic and heterotrophic growth in acetogens is not trivial (Aklujkar et al., 2017; Marcellin et al., 2016; Nagarajan et al., 2013; Tan et al., 2013). We thus aimed to determine TSSs and transcriptional features of promoter motifs and transcriptional regulators associated with essential genes (including genes of the WLP) in the model-acetogen *C. autoethanogenum*.

Our study revealed a new promoter motif and the identification of two proteins activating gene expression from the new motif (the TetR-family protein (CAETHG_0459) and the housekeeping σ^A^ (CAETHG_2917)). An alternative TetR transcriptional regulator has been previously found to be a σ factor in *Clostridium tetani*, and its homologues, TcdR in *C. difficile*, BotR in *C. botulinum*, and UviA in *C. perfringens* have also been found to regulate toxin production (Dupuy et al., 2006; Dupuy and Matamouros, 2006; Raffestin et al., 2005). In combination, these results suggest that TetR proteins can play an important role in transcriptional regulation in *clostridia*. These studies support our PPI assay potentially suggesting that the TetR-family protein might function as a σ factor in *C. autoethanogenum*, but further studies (*in vitro* transcription assay) are needed to confirm this. In fact, unequivocal demonstration of σ factor activity requires that a protein is necessary and sufficient for activation of promoter recognition and transcription initiation by RNAP, independent of any other σ factor subunit. Thus our results do not exclude the possibility that a native σ factor of the in vivo expression host (*E. coli*), e.g. σ^70^, could have induced the TetR-family protein to drive transcription from P_cauto_. Additional studies should also be performed to study whether both the σ^A^ and the TetR-family protein show an overlap in the promoter motif for transcriptional activation (Cho et al., 2014).

Notably, there are several TetR-family proteins, commonly regarded as transcriptional regulators (Cuthbertson and Nodwell, 2013), annotated in the *C. autoethanogenum* genome. In pathogenic *clostridia* these TetR-family proteins are often described as alternative σ factors, belonging to a class of σ factors called extracytoplasmic function (ECF) σ factors (Feklistov et al., 2014; Sineva et al., 2017). Their discovery led to a novel class of σ factors (group 5), which show a −35 and/or −10 conserved region in their target promoters (Dupuy et al., 2005, 2006; Dupuy and Matamouros, 2006; Staroń et al., 2009). It will be interesting to see whether transcription from P_cauto_ described here with an interspaced repetition of (A/T)G notably distinct from the canonical −35/-10 conserved regions is also activated by a novel σ factor.

Our work also shows that the housekeeping σ factor (σ^A^) in *clostridia* can activate transcription from P_cauto_ associated with essential genes for autotropic growth in acetogens. Interestingly, in another acetogen *E. limosum*, the promoter regions of genes of the WLP, hydrogenases, and ATPase contain the well-known −35 TTGACA and −10 TATAAT motifs for the housekeeping σ factor (σ^A^) (Burgess and Anthony, 2001; Song et al., 2017). This potentially indicates that the housekeeping σ^A^ in acetogens can initiate transcription from different promoter motifs and illustrates well the great extent of genetic diversity among the non-taxonomic group of acetogens. While the WLP itself is highly conserved, it is not surprising that transcriptional regulation is diverse (Drake et al., 2006; Shin et al., 2016). The work presented here also highlights the importance of P_cauto_ in other industrially relevant acetogens (Figure 2C). We believe, however, that more studies are needed for the experimental determination of TSSs and transcriptional features to facilitate a broader understanding of transcriptional regulation in acetogens.

Our findings have the potential to significantly advance the understanding of transcriptional regulation and metabolic engineering of the ancient metabolism of acetogens. Firstly, acetogen metabolism, which operates at the thermodynamic edge of feasibility (Schuchmann and Müller, 2014), seems to be wired for utilising less energy-consuming mechanisms (i.e. transcriptional vs. translational regulation) for operating under different conditions evidenced by the complexity of the condition-specific transcriptional architecture (Valgepea et al., 2018). More importantly, the discovery of P_cauto_ and a key positive transcription factor (TetR-family protein) in acetogens can lead to the mechanistic description of transcriptional regulation of arguably the first biochemical pathway on Earth (Fuchs, 2011; Russell and Martin, 2004; Weiss et al., 2016). In addition to expanding the fundamental understanding of a model acetogen, knowledge of the features controlling the expression of essential genes in acetogens could also contribute for the improvement of commercial gas fermentation for the sustainable production of fuels and chemicals. Increasing or modulation of the activity of the described TetR transcription factor (either through over-expression and/or protein engineering or by deleting transcriptional repressor genes) could enhance the uptake of C_1_ substrates through the WLP and thus improve growth and/or product formation (possibly by introducing P_cauto_ in front of key genes). It could also be used as an orthologous system in other organisms, as, for instance, the TcdR system has been used in other *Clostridium* species (Minton et al., 2016; Zhang et al., 2015). Importantly, the newly discovered promoter P_cauto_ could be harnessed to couple expression of heterologous pathways to mimic those of key central metabolism enzymes, potentially alleviating the common problem of imbalanced flux throughput between heterologous and native metabolic pathways.

## Author contributions

(i) RL, KV, MK, RT, LN and EM designed the study and the experiments (ii) RL, RG, RP, KV, CB performed the experiments. RL, KV, RT, RP, MK, SS, LN and EM analysed and interpreted the data; (iii) RL, KV, RT and EM wrote the manuscript. All authors reviewed the manuscript.

## Acknowledgements

The authors acknowledge support from the Queensland node of Metabolomics Australia (MA) at The University of Queensland, an NCRIS initiative under Bioplatforms Australia Pty Ltd. We thank Dr Christopher Howard for his helpful advice in the pull down/affinity chromatography assay and protein purification. We thank the following investors in LanzaTech’s technology: BASF, CICC Growth Capital Fund I, CITIC Capital, Indian Oil Company, K1W1, Khosla Ventures, the Malaysian Life Sciences, Capital Fund, L. P., Mitsui, the New Zealand Superannuation Fund, Petronas Technology Ventures, Primetals, Qiming Venture Partners, Softbank China, and Suncor.

## Datasets are in a publicly accessible repository

dRNA-Seq data have been deposited in the NCBI Gene Expression Omnibus depository under accession number GSE108700.

Re-annotation of *C. autoethanogenum genome* was deposited in the NCBI GenBank Third Party Annotation database under accession number BK010482.

