## Supplementary material for "A TetR-family protein activates transcription from a new promoter motif associated with essential genes for autotrophic growth in acetogens": SI1

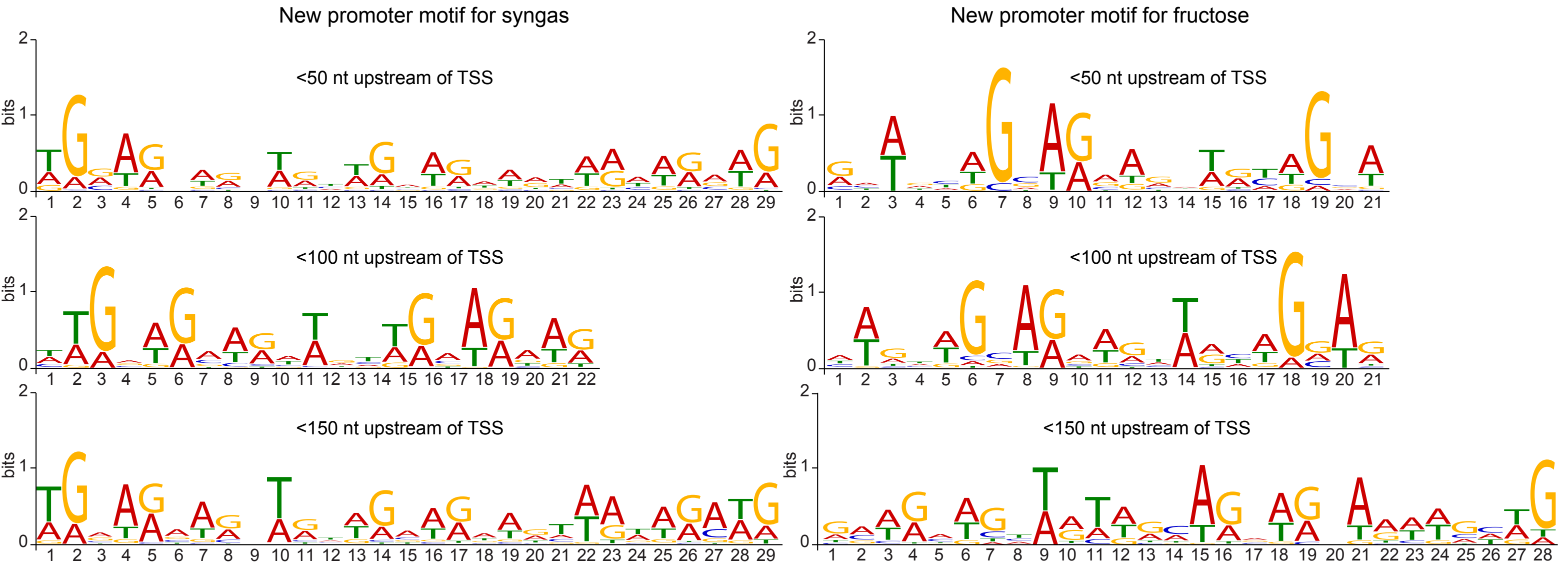


FIG S1 The new promoter motif is conserved within 50 nt upstream of TSSs. The three motifs for each condition were obtained by searching for consensus sequence motifs 50, 100, or 150 nt upstream of TSSs assigned with the new promoter motif using the MEME software (Bailey *et al*, 2009). Height of the letter indicates its relative frequency at the given position within the motif.
