## Supplementary material for "A TetR-family protein activates transcription from a new promoter motif associated with essential genes for autotrophic growth in acetogens": SI2

| **Supplementary File 2.** Molecular cloning. Bacterial strain, plasmids and primers used* | | |
| --- | --- | --- |
| **Bacterial strain** | **Description** | **Reference** |
| *Escherichia coli* BL21 | Expression host | Bioline |
| *Escherichia coli* DH5α | Cloning host | Bioline |
| **Plasmid** | **Description** | **Reference** |
| pBR322 | Cloning vector | Life Technologies |
| pACYC184 | Expression vector | NEB |
| pET28a+ | Expression vector | AddGene |
| pBR_PprpR-GFPUV | pBR322 derived plasmid built in house. GFP-uv expression plasmid under the control of prpR promoter | Lab collection |
| pET_SelT | pET28a+ derived plasmid. KanR, *T7-SelT* | This work |
| pET_TetR1^a^ | pET28a+ derived plasmid. KanR, *T7-TetR1* | This work |
| pET_TetR2 ^b^ | pET28a+ derived plasmid. KanR, *T7-TetR2* | This work |
| pET_GntR | pET28a+ derived plasmid. KanR, *T7-GntR* | This work |
| pET_Sigma70 | pET28a+ derived plasmid. KanR, *T7-Sigma70* | This work |
| pBR_ P_cauto_ _GFP | pBR322 derived plasmid. AmpR, *P_cauto_-gfpUV* | This work |
| pAC_P_cauto__GFP | pACYC184 derived plasmid. CmR, *P_cauto_-gfpUV* | This work |
| pAC_P_cauto_ 130_gfp | pACYC184 derived plasmid. CmR, 130 bp region *P_cauto_*-gfpUV | This work |
| pAC_P_cauto_ 30A_gfp | pACYC184 derived plasmid. CmR, 30 bp region *P_cauto_*-gfpUV | This work |
| pAC_P_cauto_ 30A_gfp | pACYC184 derived plasmid. CmR, 30 bp mutated region *P_cauto_*-gfpUV | This work |
| **Primer name** | **Sequence** | **Target region** |
| SelT_FWD | ctggtgccgcgcggcagccatATGGATAAAAAACAATTATTAAGAAAAC | SelT (CAETHG_2839) |
| SelT_REV | ggtgctcgagtgcggccgcaagcttattaCTAAAACTCTGTAAAAGCATCTACC |  |
| TetR2_FWD ^b^ | ctggtgccgcgcggcagccatATGGCTCAAATAAAAAAAGAC | TetR2 (CAETHG_0936) |
| TetR2_REV ^b^ | ggtgctcgagtgcggccgcaagcttattaTTACTTAGTGTATTCTGTAAGTAATCTTTTAAATC |  |
| GntR_FWD | ctggtgccgcgcggcagccatATGTCATCGCAAAATGTTAC | GntR (CAETHG_3915) |
| GntR_REV | ggtgctcgagtgcggccgcaagcttattaCTAATTATTATCTCCATAAAGAAATTTTTTG |  |
| CautoPnew_FWD | ttgacagcttatcatcgataagcttAATTAGGCGTAAATGTAAGATTATC | P_cauto_ |
| CautoPnew_REV | ctttactcatATTGTTTCCTCCTAAATGTTTTTAG |  |
| Cauto_gfp_FWD | aggaaacaatATGAGTAAAGGAGAAGAACTTTTC | gfp-UV |
| Cauto_gfp_REV | gtgataaactaccgcattaaagcttattaTTATTTGTAGAGCTCATCCATG |  |
| pET_conf(FWD) | GATATAGGCGCCAGCAACC |  |
| pET_conf(REV) | AGCCAACTCAGCTTCCTTTC |  |
| P_cauto__GFP_conf(FWD-1) | TGTCTCATGAGCGGATACATATT |  |
| P_cauto__GFP_conf(REV-1) | GTCTTGTAGTTCCCGTCATCTT |  |
| P_cauto__GFP_conf(FWD-2) | GGAAACATTCTCGGACACAAAC |  |
| P_cauto__GFP_conf(REV-2) | ATACCCACGCCGAAACAA |  |
| P_cauto__GFP_conf(FWD-3) | GGAAACATTCTCGGACACAAAC |  |
| P_cauto__GFP_conf(REV-3) | TTGTTTCGGCGTGGGTAT |  |
| * Upper cases indicate the region homologous to the target region during the PCR amplification. Lower cases indicate homologous region for Gibson assembly.  ^a^ TetR 1: CAETHG_0459  ^b^ TetR 2: CAETHG_0936 | | |
